# Metabolic plasticity facilitates a high-latitude life of feast and famine in Arctic char

**DOI:** 10.64898/2026.02.01.703141

**Authors:** Matthew J.H. Gilbert, Ella K. Middleton, Les N. Harris, Emily P. Williams, Thomas Landry, Abby R. Christopher, Simon G. Lamarre, Jean-Sébastien Moore, Ben Speers-Roesch

## Abstract

1. High northern latitudes experience extreme winters and pronounced environmental variation. Arctic animals possess remarkable and diverse physiological mechanisms to thrive under this variation, but such mechanisms are poorly studied, particularly in the aquatic realm.
2. We combined Arctic field and laboratory experiments to reveal how one of Earth’s most northerly distributed fish species, the Arctic char, seasonally adjusts its metabolism to conserve energy over winter while not feeding and to capitalize on more optimal growth conditions in summer.
3. Laboratory reared Arctic char were fed or food deprived at common summer (8°C) and winter (2°C) temperatures for >120 days. Oxygen consumption rates were assessed as a proxy for rate of aerobic metabolic energy expenditure. Cooling from 8 to 2°C reduced resting aerobic energy expenditure by 35%, while food deprivation independently drove a reduction of 46-50%. Combined, resting energy expenditure was 197% higher under simulated summer (fed at 8°C) than winter conditions (food deprived at 2°C).
4. Reductions in resting energy expenditure with food deprivation were driven, at least in part, by marked tissue-specific decreases in organ size and protein synthesis rates in tissues with central roles in growth and digestion.
5. Wild adult anadromous Arctic char exhibited, seasonal plasticity consistent with our laboratory experiments. Resting energy expenditure at 10°C was 27% higher after summer feeding than in the spring after winter fasting, while liver and digestive organ masses were also markedly reduced overwinter.
6. In contrast, peak aerobic metabolic performance and relative heart mass were largely maintained during food deprivation indicating a protection of systems that support aerobic metabolic capacity.
7. Despite recruiting strategies for substantial energy savings, wild and laboratory reared Arctic char still largely depleted their on-board fat stores with food-deprivation, underscoring that these strategies are essential when food is limited. Given that the Arctic is among the most rapidly changing regions on Earth, temperatures, productivity, and the timing of key seasonal event are all shifting with implications for the efficacy of such strategies for high-latitude animals.

## Introduction

Many animals have evolved remarkable physiological and life-history strategies to thrive in the face of temporal variation in temperature and food availability (McNab 2002, Secor and Carey 2016). In particular, at high northern latitudes (>60°N) animals face extreme winter conditions and strong environmental seasonality which can necessitate physiological plasticity, dictate life histories, and greatly restrict species’ abundance and distributions (Chown et al., 2004; Jørgensen & Johnsen, 2014) Ghalambor et al. 2006, Shuter et al. 2012, Studd et al. 2021). Yet, the physiological mechanisms underlying classic responses to winter – avoidance through migration, prolonged dormancy, or continued activity at cost – remain largely understudied at high latitudes (Marchand, 2014). Most eco-physiological research on northern animals has focused on select terrestrial mammals (e.g., hibernating rodents and bears), insects, and depauperate herpetofauna (e.g., Larson et al. 2014). Less attention has been given to understanding how the physiology of northern aquatic fauna is shaped by and facilitates life under strong environmental seasonality, including extreme winter conditions.

During winter, northern aquatic animals experience extremely low water temperatures (e.g. 0-2°C in freshwater; -1.5°C in seawater), short or non-existent photoperiods, and prolonged ice coverage (∼9-10 months year^-1^) (Hammer et al., 2022; Jørgensen & Johnsen, 2014; Mulder et al., 2018). In contrast, summer commonly involves long photoperiods, and strong pulses of productivity, while water temperatures can be unexpectedly warm, particularly in shallow areas (e.g., >20°C in a fish bearing river; Gilbert and Tierney 2018). These conditions create dramatic seasonal inconsistencies in foraging ability and food availability (Gross et al., 1988). Thus, like their terrestrial counterparts, northern aquatic animals often require energy-saving strategies to cope with prolonged winter food restriction. In temperate fishes, strategies to cope with food scarcity include voluntary fasting, inactivity, and active reductions in resting metabolism (Auer, Salin, Anderson, et al., 2016; Secor & Carey, 2016; Speers-Roesch et al., 2018; Van Leeuwen et al., 2012; Reeve et al. 2022; Middleton et al. 2024). Independent from food availability, winter cold can directly induce fasting, reduce activity, and passively slow metabolism, thereby reducing energy expenditure (Reeve et al., 2022). The underlying mechanisms and roles that these strategies play in the winter energy conservation of high-latitude fishes are not well documented. This knowledge gap is significant given that high-latitudes are warming several times faster than the average global rate (Rantanen et al., 2022). This warming has pronounced impacts on aquatic habitats (Gillis et al. 2024) and directly increases fish energy expenditure (Clarke & Johnston, 1999). Associated increases in environmental variability (Morash et al., 2018) also affects the predictability of temperatures and food availability. Together, warming, phenological shifts, and increasing environmental variability are likely to impact the efficacy of the energy management strategies that northern fishes use to navigate protracted winters and brief summers (Shuter et al., 2012; Sutton et al., 2021).

As Earth’s most northerly distributed freshwater or anadromous fish, the Arctic char (*Salvelinus alpinus*) is an intriguing aquatic species in which to study mechanisms of energy management in the context of strong seasonality and extreme winters (Klemetsen, 2010; Walseng et al., 2018). Anadromous Arctic char migrate from freshwater habitats to the more productive marine environment to feed for ∼four to six weeks each summer (Johnson, 1980; Klemetsen et al., 2003) (Harris et al. 2022). However, the elevated salinity and frigid (-1.5°C) winter temperatures of the Arctic Ocean drive Arctic char to return to more tolerable conditions in freshwater each fall. Once back in freshwater, Arctic char become largely inactive and generally cease feeding over winter (∼10 mo. yr^-1^) (Boivin & Power, 1990; Dutil, 1986; Jørgensen & Johnsen, 2014). During this prolonged food deprivation they commonly lose ∼30% of their total energy, and even more in years that they spawn (Dutil, 1986). Thus, the annual energy budgets of adult anadromous Arctic char can be entirely dependent on brief but rapid energy acquisition over summer and energy conservation over the protracted winter (Harwood & Babaluk, 2014; Johnson, 1980), paralleling many northern terrestrial species (e.g. grizzly bears (Schindler et al. 2013).

Here, our principal aim was to elucidate mechanisms for energy management in fishes under extreme winter conditions and to explore trade-offs for metabolic performance using Arctic char as our model species. We hypothesized that, like iconic mammalian hibernators (e.g., bears; Tøien et al. 2011), overwintering Arctic char conserve energy through separate mechanisms independently associated with cold and food deprivation. Specifically, we hypothesized that Arctic char undergo a passive slowing of resting metabolism with cooling, and an active suppression of resting metabolism when food-deprived. To thoroughly test this hypothesis, we took the rare approach of combining detailed physiological experiments in a laboratory and in an Arctic field setting. This approach allowed us to test the effects of specific environmental factors in a controlled environment and then reveal how those effects manifest in nature (Turko et al. 2023). In the laboratory, we fed or starved Arctic char for >4 months at common winter (2°C) or summer (8°C) water temperatures. We performed organismal (resting oxygen consumption and aerobic metabolic performance) and sub-organismal (*in vivo* protein synthesis rates, organ sizes, and energy storage) assessments for an integrative understanding of energy management across levels of biological organization (Somero, 2015). In the field, we made complementary assessments in wild anadromous Arctic char captured at the end of winter (at the start of re-feeding) and at the end of summer (after summer feeding). Beyond typical temperature effects on aerobic metabolism, we predicted that resting oxygen consumption would be reduced following food deprivation in association with tissue-specific atrophy and reductions in protein synthesis in tissues associated with growth and digestion. Given the tissue-specific selectivity of these predicted changes we expected peak oxygen consumption and capacity to be largely maintained. Combined, these field and laboratory approaches are designed to advance our fundamental mechanistic understanding of how high-latitude fishes navigate extreme seasonal variation with implications for a rapidly changing north.

## Methods

### Experiment 1: The effects of temperature and prolonged food-deprivation on the energetics of Arctic char in a laboratory setting

#### Experimental animals

Arctic char (mass: 133.1 ± 38.7 g, fork length: 245.1 ± 23.8 mm; mean ± SD) were obtained in December 2020 from Aquaculture Gaspésie (Gaspé, QC, Canada) at a size greater than that typical at first migration (∼50 g, ∼190 mm; Gilbert et al., 2016). This aquaculture strain of Arctic char originated from a wild anadromous population at Nauyuk Lake, NU, Canada, <100km from our field study site. Fish were transported in aerated freshwater at ∼ 4°C to UNB where they were housed in 90 L glass aquaria (3-5 fish per aquarium) at 8°C ± 0.5°C and fed daily a ∼0.5% body mass ration (23 kJ g^-1^; 40A 4.0 mm BioTrout, Bio-Oregon, Washington State, USA) for four weeks before starting the experiment. The fish were held under a photoperiod (10L:14D) characteristic of early or late winter in the Canadian High Arctic, with a simulated sunrise and sunset (30 min each) for the entire duration of holding and the experiment.

#### Experimental system and design

The aquaria were partially submerged in four aerated ∼400L fibreglass troughs (four aquaria per trough) containing filtered, dechlorinated municipal freshwater with a slow continuous inflow of freshwater to further maintain water quality. Water was continuously recirculated to the aquaria from the surrounding trough and aquaria in each trough were split evenly into fed and starved treatments (detailed below). Two troughs were maintained at 8°C ± 0.5°C and two were cooled to from 8°C to 2°C ± 0.5°C over ∼10 hours and then held at 2°C for ∼24 hours before initial respirometry measurements at day ‘0’; subsequent measurements were considered relative to this initial measurement as previously described (Middleton et al. 2024). Within each temperature treatment, fish were either fed daily (∼0.5% body mass ration) or starved, resulting in four treatments: fed at 8℃, starved at 8℃, starved at 2℃ and fed at 2°C. Importantly, for the first 60 days of the experiment several fed fish, particularly at 2℃, had a low appetite which is common over winter in Arctic char given their natural winter biology (Tveiten et al., 1996). This variable appetite increased variation in growth rates among individuals which allowed for additional correlative analyses of metabolic parameters as a function of growth rate (Figure S1a). Experiments ended in spring at which point all but two fish in the fed treatments had resumed feeding and between respirometry measurements made on day 60 and 120 allowing for robust comparisons of energetics between fed and starved fish as intended (Figure S1b).

#### Measurements of oxygen consumption rate (ṀO_2_) and spontaneous activity

Automated intermittent-closed oxygen respirometry was used to assess oxygen consumption rates (*Ṁ*O_2_; mgO_2_ kg^-1^ h^-1^) to estimate aerobic metabolic rates (Svendsen et al. 2016) exactly as described by Middleton et al. (2024). Respirometry lasted 48 hours. Following each respirometry trial, fish were weighed and lengthed, and Fulton’s condition factor (K; see Data Analysis) was calculated. SMR (*Ṁ*O_2standard_) was estimated at 0, 30, 60 and 120 days as the 20^th^ percentile of all *Ṁ*O_2_ measurements in the final 24 hours of the 48-hour measurement period (Chabot et al., 2016) as in Middleton et al. (2024). We ensured that the 20^th^ percentile method represented resting oxygen consumption by simultaneously recording the spontaneous activity of fish in the respirometers to confirm inactivity in all treatment groups (see below) and by estimating *Ṁ*O_2standard_ at zero activity (see Supplemental Information; Table S1).

The maximum oxygen consumption rate was estimated (*Ṁ*O_2max_) exactly as in Middleton et al. (2024) at the final time point (120 days) only, immediately following the *Ṁ*O_2standard_ measurement period. The difference between *Ṁ*O_2standard_ and *Ṁ*O_2max_ was calculated as the absolute aerobic scope (AAS) of each fish. Spontaneous activity of each fish in the respirometers at the final sampling point (120 days) was measured simultaneously with resting *Ṁ*O_2_ from video using an automated tracking software (Ethovision XT v.16, Noldus Information Technology BV, Netherlands) as previously described (Middleton et al. 2024).

#### Measurements of tissue energetics

Following the final respirometry sampling event (120 days), fish were allowed to recover under their treatment conditions for at least 5 days before chemical euthanasia (300 mg L^-1^ MS-222 and 600 mg L^-1^ sodium bicarbonate, Sigma-Aldrich, MO, USA) and immediate assessment of body fat, protein synthesis rates, hematocrit, blood glucose, and organ masses as previously described (Middleton et al. 2024)(Cassidy et al., 2016; Lamarre et al., 2015). Tissue samples were manually ground on liquid nitrogen with mortar and pestle and stored, along with plasma samples, at -80°C until analysis. The remaining carcass was ground and freeze dried for 48 hours (-50°C and 0.12 mBar), followed by homogenization of the dry tissue. Energy density (kJ g dry mass^-1^) of the carcass was determined by bomb calorimetry (6765 Combination Calorimeter, Parr Instrument Company, IL, USA). A commercial assay kit (10010303, Cayman Chemical, MI, USA) was used to quantify the triglyceride (TG) contents of the plasma (Frøiland et al., 2012), gut and white muscle. The Bradford assay was used to quantify protein content in the liver, gut, white muscle and carcass following protocols detailed in Middleton et al. (2024).

### Experiment 2: Examining energy savings over winter in wild anadromous Arctic char

#### Experimental animals

The field study took place at Palik (Byron Bay), in the Kitikmeot region of Nunavut, in the central Canadian Arctic (68°94’N, 108°52’W; Figure S2). In early July 2019 and late August 2022, wild anadromous Arctic char (Table 3) were collected at the mouth of the Lauchlan River using a continuously monitored 139 mm mesh gill net or by angling with barbless hooks. Fish caught in July had just migrated to the ocean at the end of winter and had been fasting for ∼10 months, whereas fish caught in August at the end of summer would have fed at sea for 4-6 weeks. Following capture, Arctic char were held for 24-48 hours in mesh pens (∼1m x 1m x1m) submerged in the estuary near the site of capture to recover and evacuate their gut before respirometry. Water temperatures at the capture site and during holding ranged from 8.4°C-8.8°C in July 2019 and 6.5°C-10.6°C in August 2022. Given that some samples were collected opportunistically and that we had refined the sampling protocol in 2022, there were variations in organ availability between years and seasons (Table S3). Three ventricle measurements, and correspondingly, three gut:heart measurements were identified as outliers using the 1.5x IQR method (Tukey, 1977) and removed from analysis as the value were not biologically realistic.

#### Measurement of oxygen consumption rate (ṀO_2_)

Automated intermittent-flow respirometry was conducted in a canvas wall-tent (2.4×3.0 m) with power supplied by a mobile research laboratory (Arctic Research Foundation, Winnipeg, Canada) described by Gilbert et al (2020). We used two cylindrical 40.7 L acrylic respirometers (24 cm internal diameter × 90 cm long) (Loligo Systems, Viborg, Denmark) which were housed in a 250 L plastic lined PVC frame (42×30×12 inches) in 2019 and a circular 615 L polypropylene bath (63” diameter × 12” height) in 2022. Temperature was maintained at 10°C ± 1°C in both years. Respirometers were fitted with the same optodes as above. The total intermittent flow cycle duration was 10 minutes with respirometers flushed for two minutes and closed for eight.

To start, fish were individually chased to exhaustion (2019: 3.2 ± 0.5 min; 2022: 3.8 ± 0.7 min), which was quantified as the point when the fish did not respond to a caudal or mid-body grab for more than five seconds (Gilbert, 2020), and then exposed to air for ∼1 min (2019: 1.10 ± 0.2 min; 2022: 1.00 ± 0.1 min). Fish were then rapidly sealed in the respirometers and a manual intermittent flush procedure was repeated twice to obtain slope estimates for *Ṁ*O_2max_ to maximize our closed duration during *Ṁ*O_2max_ assessment without allowing DO to drop below 75% air saturation. Fish were then returned to an automated intermittent flush cycle and held in the respirometers for ∼20 hours. The first six hours of recordings were excluded as an adjustment and recovery period and were removed from the estimates of resting oxygen consumption (*Ṁ*O_2rest_). Given that these field experiments were necessarily shorter than ideal for estimates of *Ṁ*O_2standard_ and that we were working with wild fish in confinement, we use the term *Ṁ*O_2rest_ instead of *Ṁ*O_2standard_ to represent resting metabolic demands for this experiment. Regardless, oxygen consumption had declined to a low, stable level during the measurement period (Figure 4a). The analysis of *Ṁ*O_2rest_ and *Ṁ*O_2max_ was done as described for 8°C fish in Experiment 1 but using a slope duration of five minutes and with the exception that measurements for *Ṁ*O_2rest_ were taken as the lowest 20^th^ percentile of the data recorded over a 12-hour measurement period. AAS was calculated as above. Background respiration was accounted for using 10-minute closed periods after fish were removed following each respirometry trial. To minimize the microbial load and accumulation of wastes, water was changed between each trial.

#### Tissue collection and analysis

Organ samples were taken from fish after respirometry measurements and opportunistically from fish harvested for subsistence purposes or as part of other ecological studies at the same time as respirometry was conducted. Fish were euthanized by cervical percussion immediately following capture or after respirometry trials. Tissue samples (ventricle, liver, gut and white muscle) were removed, separated, weighed and frozen at -20°C until analysis. All tissue samples were reweighed at UNB in 2022 and 2023 and freeze dried (FreeZone 12 L, Labconco Corp., MO, USA) for 48 hours (-50°C and 0.12 mBar) to obtain dry weights; the only exception were the ventricles which were only weighed in the field. Bomb calorimetry and TG and protein assays were conducted as in Experiment 1 to obtain the energy density of white muscle and TG and protein contents of the gut and white muscle.

### Data analysis

Fulton’s condition factor (K) was calculated using the following equation:

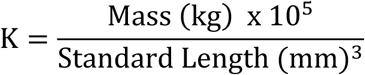

The thermal sensitivity quotient (Q_10_) was calculated using the following equation:

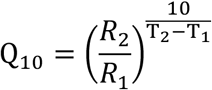

Where R_1_ was the rate for a biological process at the lower temperature, T_1_, and R_2_ is the rate at the higher temperature, T_2_.

To calculate size-corrected organ masses in Experiment 1, each organ mass or length were modelled in separate models. Size-adjusted organ mass or length was obtained by adding the model residuals to the model estimate at the average body mass (0.162 kg) or length (261 mm; intestine length only). To generate a somatic index (%), the adjusted organ masses were divided by the average body mass or length and multiplied by 100. This method accounts for any deviation from the assumption of a 1:1 mass or length scaling relationship. For Experiment 2, somatic indices were calculated using the same method, except they were centered to the average body length in a similar manner to Armstrong & Bond (2013). Dry organ mass was used for all organs except the ventricle. For both experiments, we calculated the gut:heart ratio by dividing wet gut mass by wet ventricular mass with the goal of examining how the heart and digestive organs changed relative to one another among treatments.

#### Statistical analysis

Statistical analyses were performed using R Studio (Core Team, 2014) and Prism v.9 (GraphPad Software, CA, USA) and data visualization was completed using Prism v.9. Statistical significance was set at p<0.05.

Linear models (LM) or linear mixed effects models (LMM) were used to analyze the data unless the assumptions of normality and homoscedasticity of the residuals were not met.

Assumptions were tested using diagnostic plots of residuals and Shapiro-wilks tests for normality. If assumptions were not met, generalized linear models or generalized linear mixed effects models (GLM or GLMM; family=Gamma, link=log) were used. For Experiment 1, models were developed with each metric as a function of temperature, feeding status and time (if applicable). Body mass or body length were included as covariates when applicable (see Tables S4-S6) to account for allometric scaling. Fish ID was included as a random factor for models with repeated measures. For Experiment 2, each metric was assessed as a function of time of year (end of winter or end of summer), with body mass included as a covariate for analysis of *Ṁ*O_2_ and body length for somatic indices as in Armstrong and Bond (2013).

Type II Wald chi-square tests were performed on all models using the “car” package (Fox et al., 2007) followed by pairwise comparisons of estimated marginal means (“emmeans” package; Lenth et al., 2019) with a Holm adjustment for multiple comparisons.

In data presentations, the allometric scaling of *Ṁ*O_2_ with body mass was accounted for by adjusting *Ṁ*O_2standard_ to the average body mass across all treatment groups; we added the residuals from the mass vs. *Ṁ*O_2_ relationship in each model to the predicted value at the average mass (0.144 kg) to obtain a mass centered *Ṁ*O_2_ value, which was then divided by the average mass to generate mass-specific values for presentation. The laboratory 120-day metabolic performance data was adjusted in the same way, except that *Ṁ*O_2_ was adjusted to an average body mass of 0.161 kg because two individuals were removed from the fed at 2°C treatment group due to lack of feeding. For the field study, the allometric scaling of *Ṁ*O_2_ was accounted for in data presentation as described above but with an adjustment to the average body mass of 4.21 kg.

## Results

### Experiment 1: The effects of cold and prolonged starvation on the energetics of Arctic char in a laboratory setting

#### Growth metrics

The body length, mass and K increased throughout the experiment in fed fish, with the minor exception of a small loss of mass and thus, lower K, between the first and second measurements in fed fish at 2°C because some individuals slowed or paused feeding (Table 1, Table S4). In starved fish, mass and K generally decreased over time, while length did not change (Table 1, Table S4).

**Table 1.**
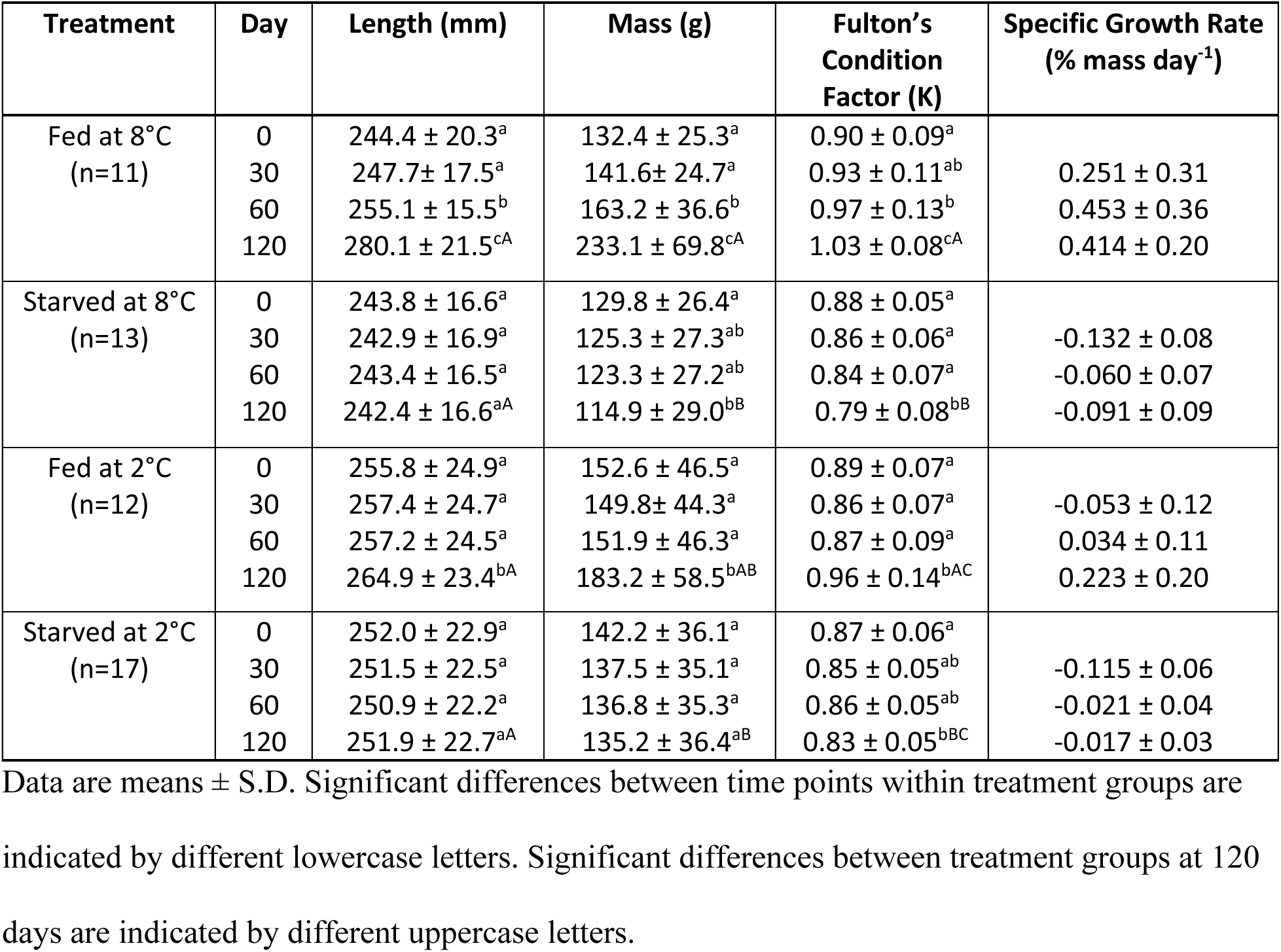
Morphometrics and specific growth rates of juvenile Arctic char that were fed daily (0.5% body mass ration) or starved at 8°C or 2°C for 120 days. Data is only included for fish that had complete datasets (i.e., mass and length measured at every time point).

| Treatment | Day | Length (mm) | Mass (g) | Fulton's Condition Factor (K) | Specific Growth Rate (% mass day<sup>-1</sup>) |
| --- | --- | --- | --- | --- | --- |
| Fed at 8°C<br>(n=11) | 0 | 244.4 ± 20.3<sup>a</sup> | 132.4 ± 25.3<sup>a</sup> | 0.90 ± 0.09<sup>a</sup> |  |
|  | 30 | 247.7 ± 17.5<sup>a</sup> | 141.6 ± 24.7<sup>a</sup> | 0.93 ± 0.11<sup>ab</sup> | 0.251 ± 0.31 |
|  | 60 | 255.1 ± 15.5<sup>b</sup> | 163.2 ± 36.6<sup>b</sup> | 0.97 ± 0.13<sup>b</sup> | 0.453 ± 0.36 |
|  | 120 | 280.1 ± 21.5<sup>cA</sup> | 233.1 ± 69.8<sup>cA</sup> | 1.03 ± 0.08<sup>cA</sup> | 0.414 ± 0.20 |
| Starved at 8°C<br>(n=13) | 0 | 243.8 ± 16.6<sup>a</sup> | 129.8 ± 26.4<sup>a</sup> | 0.88 ± 0.05<sup>a</sup> |  |
|  | 30 | 242.9 ± 16.9<sup>a</sup> | 125.3 ± 27.3<sup>ab</sup> | 0.86 ± 0.06<sup>a</sup> | -0.132 ± 0.08 |
|  | 60 | 243.4 ± 16.5<sup>a</sup> | 123.3 ± 27.2<sup>ab</sup> | 0.84 ± 0.07<sup>a</sup> | -0.060 ± 0.07 |
|  | 120 | 242.4 ± 16.6<sup>aA</sup> | 114.9 ± 29.0<sup>bB</sup> | 0.79 ± 0.08<sup>bB</sup> | -0.091 ± 0.09 |
| Fed at 2°C<br>(n=12) | 0 | 255.8 ± 24.9<sup>a</sup> | 152.6 ± 46.5<sup>a</sup> | 0.89 ± 0.07<sup>a</sup> |  |
|  | 30 | 257.4 ± 24.7<sup>a</sup> | 149.8 ± 44.3<sup>a</sup> | 0.86 ± 0.07<sup>a</sup> | -0.053 ± 0.12 |
|  | 60 | 257.2 ± 24.5<sup>a</sup> | 151.9 ± 46.3<sup>a</sup> | 0.87 ± 0.09<sup>a</sup> | 0.034 ± 0.11 |
|  | 120 | 264.9 ± 23.4<sup>bA</sup> | 183.2 ± 58.5<sup>bAB</sup> | 0.96 ± 0.14<sup>bAC</sup> | 0.223 ± 0.20 |
| Starved at 2°C<br>(n=17) | 0 | 252.0 ± 22.9<sup>a</sup> | 142.2 ± 36.1<sup>a</sup> | 0.87 ± 0.06<sup>a</sup> |  |
|  | 30 | 251.5 ± 22.5<sup>a</sup> | 137.5 ± 35.1<sup>a</sup> | 0.85 ± 0.05<sup>ab</sup> | -0.115 ± 0.06 |
|  | 60 | 250.9 ± 22.2<sup>a</sup> | 136.8 ± 35.3<sup>a</sup> | 0.86 ± 0.05<sup>ab</sup> | -0.021 ± 0.04 |
|  | 120 | 251.9 ± 22.7<sup>aA</sup> | 135.2 ± 36.4<sup>aB</sup> | 0.83 ± 0.05<sup>bBC</sup> | -0.017 ± 0.03 |

Data are means ± S.D. Significant differences between time points within treatment groups are indicated by different lowercase letters. Significant differences between treatment groups at 120 days are indicated by different uppercase letters.

#### Oxygen consumption rate

There was a significant three-way interaction between temperature, feeding status and time on *Ṁ*O_2standard_ (*X*^2^_(3)_=10.65, p=0.014; Table S4). Following the initial cooling, *Ṁ*O_2standard_ was markedly lower at 2°C (Q_10_’s of 3.3 - 3.5) (p<0.0001 for all comparisons; Figure 1a, Table S2, Table S4). The difference in *Ṁ*O_2standard_ between the two temperatures declined with time within fed and starved treatments, resulting in lower Q_10_’s after 120 days (2.0-2.4; Table S2). The *Ṁ*O_2standard_ of fed fish at 8°C marginally increased (+14%, p=0.069) but decreased by 33% in starved fish at 8°C over 120 days (p<0.0001) (Figure 1a, Table S4). Following acclimation to 2°C for 120 days, fed fish increased *Ṁ*O_2standard_ by 40% (p<0.0001), while starved fish reduced *Ṁ*O_2standard_ by 8% (p=0.257), with this reduction occurring only between 60 and 120 days (Figure 1a, Table S4). Thus, after 120 days the *Ṁ*O_2standard_ of starved fish was markedly lower than fed fish at both temperatures (2°C: -46%; 8°C: -50%) (Figure 1b; Table S5). Furthermore, within fed groups, specific growth rate had a strong positive correlation with *Ṁ*O_2standard_ at both temperatures (2°C: R^2^=0.553, p<0.0001; 8°C: R^2^=0.565, p<0.0001) (Figure S1a).

**Figure 1.**
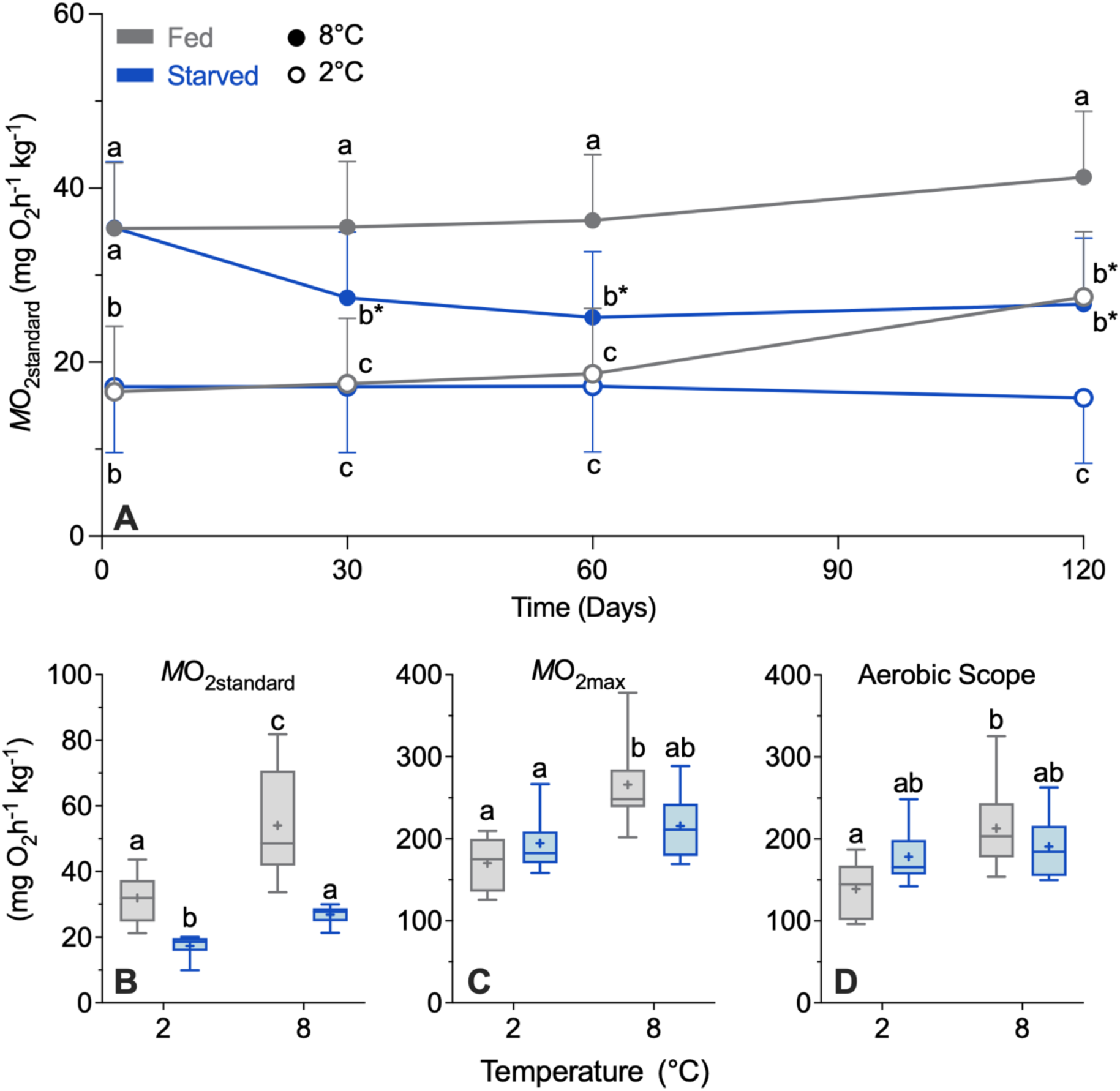
Oxygen consumption rates (*Ṁ*O_2_; respiration) and aerobic capacity of lab-reared Arctic char over 120 days of daily feeding (0.5% body mass ration) or prolonged starvation at 2°C or 8°C (Experiment 1). Data are show n for the standard rate of oxygen consumption (*Ṁ*O_2standard_; mgO_2_ h^-1^ kg^-1^) over the entirety of the experiment in all fish (A) ^where feeding^ treatments started at ‘day 0’ and temperature treatment for 2°C groups started 24 hours prior to the ‘day 0’ measurement following acute cooling (∼10 hours). Data are also shown for standard and maximum oxygen consumption (*Ṁ*O_2standard_, B; *Ṁ*O_2max_, C) and absolute aerobic scope (D) following 120 days of exposure to treatments; for fed fish, only fish that grew were included in analysis. Data are estimated marginal means (± S.E.M.) generated from GLMM and adjusted to a common body mass of 0.144 kg (A; fed at 2°C: n=10 (except for 60 days where n=8); fed at 8°C: n=10 (except for 0 days where n=11 and 120 days where n=9); starved at 2°C: n=10 (except for 120 days where n=11); starved at 8°C: n=10) or 0.161 kg (B, C, D; fed at 2°C: n=8; fed at 8°C: n=11, starved at 2°C: n=11; starved at 8°C: n=10). The boxplots represent the median and interquartile range with whiskers indicating the 5-95% percentiles; the cross represents the mean. Dissimilar letters indicate significant differences between treatment groups (within a time point; A-D), and an asterisk indicates a difference from time zero within a treatment group (A). Different letters indicate significant differences between treatment groups (B,C,D).

*Ṁ*O_2max_ was only higher in fed fish at 8°C compared with fed and starved fish at 2°C (p=0.003 and p=0.018, respectively) (Figure 1c, Table S6). As a result, AAS showed a similar pattern although AAS of fed fish at 8°C was only significantly greater compared with fed fish at 2°C (p=0.013; Figure 1d, Table S6).

#### In vivo protein synthesis rates

Ventricle, liver and gut *K_s_* decreased with temperature (ventricle: *X*^2^_(1)_=66.52, p<0.0001; liver: F_(1,36)_=44.25, p<0.0001; gut: *X*^2^_(1)_=18.68, p<0.0001) and starvation (ventricle: *X*^2^_(1)_ =43.68, p<0.0001; liver: F_(1,36)_=23.90, p<0.0001; gut: *X*^2^_(1)_=153.78, p<0.0001). Ventricle *K_s_* decreased with starvation by 37% at 2°C and 38% at 8°C (p<0.001 for both; Figure 2a, Table S6). Liver *K_s_* was lower in starved fish (27% at 2°C and 30% at 8°C) but the difference was only significant at 8°C (2°C: p=0.056 and 8°C: p<0.001; Figure 2b, Table S5). Gut *K_s_* decreased with starvation by 66% at 2°C and 68% at 8°C (p<0.0001 for both; Figure 2c, Table S6). Conversely, muscle *K_s_* was 95-155% higher in fed fish than starved fish (p<0.0001; Figure 2d, Table S6). *K_s_* had a notable positive correlation with *Ṁ*O_2standard_ in all measured tissues; the strongest correlation to *Ṁ*O_2standard_ was ventricle *K_s_* (R^2^=0.626, p<0.0001), followed by liver *K_s_* (R^2^=0.608, p<0.0001), then gut *K_s_* (R^2^=0.533, p<0.0001) and lastly, white muscle *K_s_* (R^2^=0.332, p<0.0001) (Figure 2e-h).

**Figure 2.**
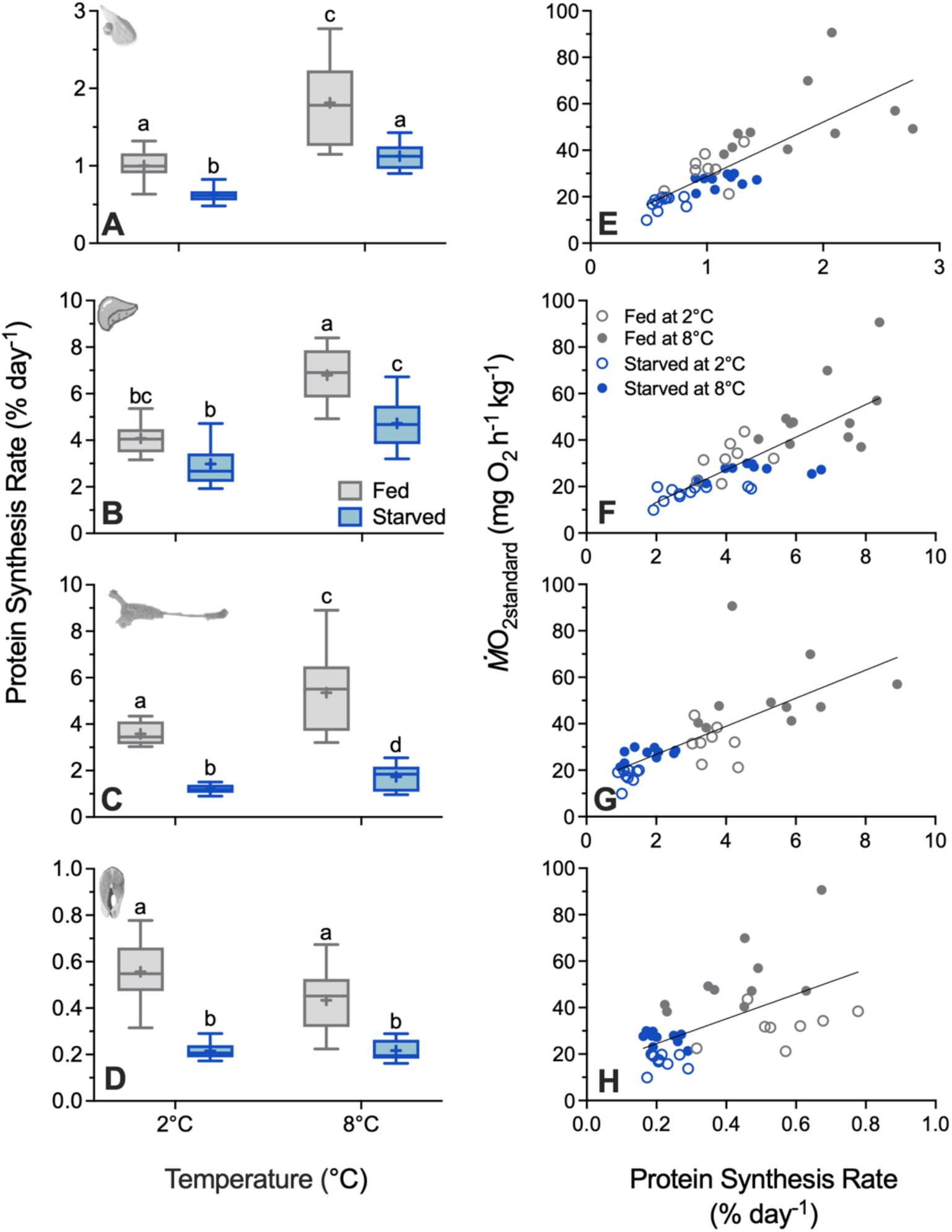
The *in vivo* fractional protein synthesis rates (*K_s_*; A-D) of juvenile Arctic char after 142 days of daily feeding (0.5% body mass ration) or prolonged starvation at 2°C or 8°C (Experiment 1) with corresponding linear regressions against *Ṁ*O_2standard_ (E-H). Data are shown for the ventricle (A&E), liver (B&F), gut (C&G), and white muscle (D&H). Boxplots represent the median and interquartile range with whiskers indicating the 5-95% percentile; the cross represents the mean (fed at 2°C; n=8, fed at 8°C; n=10 (except for liver *K_s_* where n=11), starved at 2°C; n=11 (except for gut *K_s_* where n=9 and muscle *K_s_* where n=10), starved at 8°C; n=10). Different letters represent significant differences between treatment groups. Linear regressions are shown for the (E) ventricle (R^2^=0.626, p<0.0001), (F) liver (R^2^=0. 608, p<0.0001), (G) gut (R^2^=0.533, p<0.0001) and (H) white muscle (R^2^=0. 332, p<0.0001).

#### Carcass and organ composition

The moisture content of starved fish was 4-6% higher than fed fish but the difference between fed and starved fish was only significant at 8°C (p=0.008; Table 3, Table S5). Starved fish had 38-48% less body fat than fed fish at both temperatures (2°C: p=0.001, 8°C: p<0.0001; Table 3, Table S5). Fed fish were more energy dense (+6-18%) than starved fish at both temperatures (2°C: p=0.032, 8°C: p=0.0001; Table 3, Table S5). Gut TG content and total relative gut TG content were 147-755% higher in fed fish compared with starved fish but the difference was only significant at 8°C (p=0.001, p=0.023, respectively; Table 3, Table S6). Muscle TG content was 156-246% higher in fed fish than starved fish but the difference was only significant at 8°C (p=0.006; Table 3, Table S6). There were little or no differences in protein content of the carcass and protein content or total relative protein content in the liver, gut and white muscle, except for total relative liver protein content, which was higher in fed fish than starved fish (2°C: +94%; 8°C: +63%) (see Supplemental Information for more detail; Table 3, Table S5).

#### Organ sizes

After accounting for differences in body mass, ventricle mass did not differ between treatment groups (Figure 3a, Table S5). Temperature and feeding status had interactive effects on liver and stomach mass (liver: F_(1,35)_=5.93, p=0.020; Table S5; stomach: *X*^2^_(1)_=9.60, p=0.002; Table S6) with organ mass only differing between temperatures when fish were fed. Livers (2°C: -40%; 8°C: -24%) and stomachs (2°C: -30%; 8°C: -52%) were significantly smaller in starved fish at both temperatures (Figure 3b,c). Pylorus and intestine masses were 35-52% greater in fed fish compared with starved fish (Figure 3d,e, Table S5,6). The length of the intestine was 12% greater in fed fish compared with starved fish at 2°C (p<0.0001) but was 4% smaller in fed fish compared with starved fish at 8°C (p<0.001) (Figure 3f, Table S5). Relative gut mass was greater (+48-64%) in fed fish compared with starved fish at both temperatures (p<0.01 for both; Figure 3g, Table S5) and the resulting gut:heart ratio was significantly higher in fed fish at both temperatures (p<0.0001 for both; Figure 3h, Table S6).

**Figure 3.**
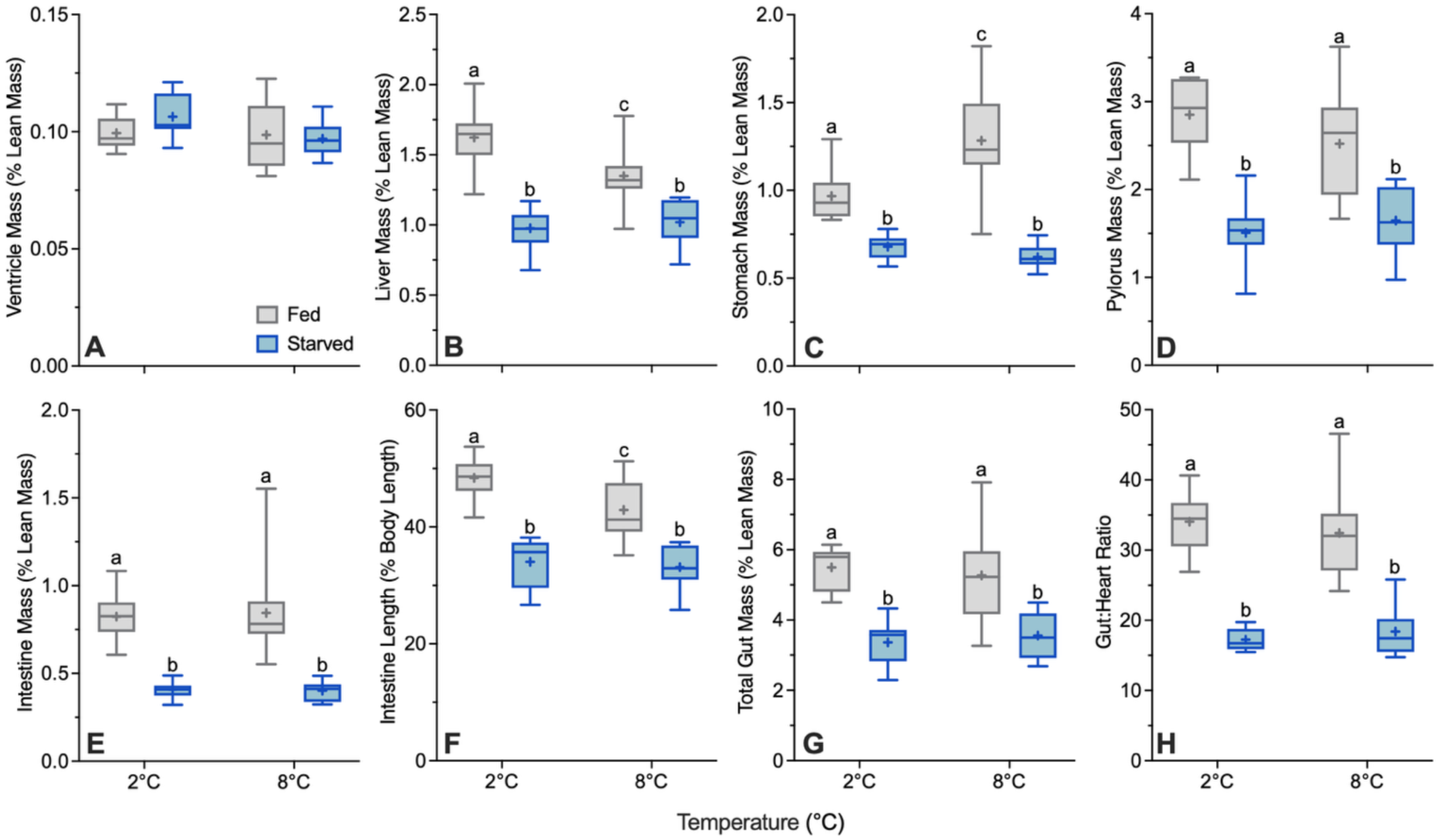
Organ size indices for laboratory reared Arctic char after 142 days of daily feeding (0.5% body mass ration) or prolonged starvation at 2°C or 8°C (Experiment 1). Organ mass as a percentage of lean body mass is shown for the (A) ventricle, (B) liver, (C) stomach, (D) pylorus, (E) intestine, and (G) total gut; (F) intestine length is shown relative to body length. Organ mass were adjusted to a common body mass of 0.162 kg to account for allometry prior to calculating relative masses. Intestine length was adjusted to a common body length of 261mm. The (H) gut to heart ratio is shown to indicate how the central organs of the digestive and circulatory systems change relative to one another. Boxplots represent the median and interquartile range with whiskers indicating the 5-95% percentile; the cross represents the mean (fed at 2°C; n=8, fed at 8°C; n=11, starved at 2°C; n=11, starved at 8°C; n=10 (except for intestine length where n=9). Different letters represent significant differences between treatment groups.

#### Blood parameters

Plasma glucose did not differ significantly among treatment groups (Table 2, Table S6).

**Table 2.** Energetic and blood parameters of juvenile Arctic char exposed to 2°C or 8°C while either fed daily (0.5% body mass ration) or starved for 142 days. Glucose was measured in plasma. Exceptions to the sample sizes are indicated below the values. Data are means ± S.E.M. Significant differences between treatment groups are indicated by different letters.

|  | **Fed at 8°C**<br>**(n=11)** | **Starved at 8°C**<br>**(n=10)** | **Fed at 2°C**<br>**(n=8)** | **Starved at 2°C**<br>**(n=11)** |
| --- | --- | --- | --- | --- |
| **Moisture Content (%)** | 72.00 ± 0.66<sup>a</sup> | 76.50 ± 1.51<sup>b</sup> | 73.06 ± 0.78<sup>ab</sup> | 76.03 ± 0.48<sup>b</sup> |
| **Body Fat (%)** | 5.89 ± 0.44<sup>a</sup> | 3.09 ± 0.19<sup>b</sup> | 5.11 ± 0.42<sup>a</sup> | 3.17 ± 0.24<sup>b</sup> |
| **Energy Density (kJ g dry mass<sup>-1</sup>)\*** | 24.67 ± 0.49<sup>a</sup><br>(n=10) | 20.90 ± 0.66<sup>b</sup> | 23.94 ± 0.25<sup>a</sup> | 21.80 ± 0.59<sup>b</sup> |
| **Carcass Protein Content (mg g<sup>-1</sup>)\*** | 314.72 ± 34.64<sup>a</sup> | 499.91 ± 63.49<sup>a</sup> | 333.49 ± 35.66<sup>a</sup> | 505.70 ± 52.61<sup>a</sup> |
| **Liver Protein Content (mg g<sup>-1</sup>)\*** | 160.33 ± 6.60<sup>a</sup> | 172.73 ± 3.28<sup>a</sup> | 173.20 ± 5.13<sup>a</sup> | 164.66 ± 5.28<sup>a</sup><br>(n=10) |
| **Total Relative Liver Protein Content (mg g body mass<sup>-1</sup>)** | 2.21 ± 0.13<sup>b</sup> | 1.36 ± 0.08<sup>c</sup> | 2.78 ± 0.16<sup>a</sup> | 1.43 ± 0.05<sup>c</sup><br>(n=10) |
| **Gut Protein Content (mg g<sup>-1</sup>)\*** | 44.81 ± 7.24<sup>a</sup><br>(n=10) | 73.17 ± 8.19<sup>ab</sup><br>(n=8) | 49.95 ± 6.33<sup>a</sup><br>(n=7) | 90.33 ± 8.30<sup>b</sup> |
| **Total Relative Gut Protein Content (mg g body mass<sup>-1</sup>)** | 2.19 ± 0.27<sup>a</sup><br>(n=10) | 1.96 ± 0.22<sup>a</sup><br>(n=8) | 2.80 ± 0.43<sup>a</sup><br>(n=7) | 2.69 ± 0.26<sup>a</sup> |
| **White Muscle Protein Content (mg g<sup>-1</sup>)\*** | 187.00 ± 8.93<sup>a</sup><br>(n=9) | 162.75 ± 9.04<sup>a</sup><br>(n=9) | 190.40 ± 6.00<sup>a</sup> | 168.32 ± 6.14<sup>a</sup><br>(n=9) |
| **Gut Triglyceride Content (mg g<sup>-1</sup>)\*** | 59.42 ± 10.72<sup>a</sup> | 14.55 ± 3.60<sup>b</sup><br>(n=9) | 33.05 ± 7.17<sup>ab</sup><br>(n=7) | 13.39 ± 4.10<sup>b</sup> |
| **Total Relative Gut Triglyceride (mg g body mass<sup>-1</sup>)** | 3.25 ± 0.75<sup>a</sup> | 0.38 ± 0.09<sup>b</sup><br>(n=9) | 1.80 ± 0.42<sup>ab</sup><br>(n=7) | 0.38 ± 0.11<sup>b</sup> |
| **White Muscle Triglyceride (mg g<sup>-1</sup>)\*** | 12.48 ± 3.51<sup>a</sup><br>(n=10) | 3.61 ± 0.50<sup>b</sup><br>(n=9) | 18.01 ± 6.38<sup>a</sup> | 7.04 ± 1.34<sup>ab</sup><br>(n=9) |
| **Plasma Glucose** | 4.56 ± 0.18<sup>a</sup> | 3.48 ± 0.08<sup>a</sup> | 5.08 ± 0.36<sup>a</sup> | 4.87 ± 0.86<sup>a</sup> |
| **Hematocrit (%)** | 34.90 ± 1.21<sup>a</sup> | 29.23 ± 1.60<sup>ab</sup> | 28.59 ± 1.22<sup>b</sup> | 30.73 ± 1.80<sup>ab</sup> |
| **Plasma Triglyceride (mg dL<sup>-1</sup>)** | 481.13 ± 36.67<sup>a</sup> | 183.12 ± 47.07<sup>c</sup> | 842.01 ± 101.64<sup>b</sup> | 210.18 ± 30.83<sup>c</sup> |

\*Contents are per unit dry mass

Hematocrit was affected by a significant interaction between temperature and feeding status (F_(1,36)_=6.48, p=0.015) that arose because temperature did not impact hematocrit unless the fish were fed (Table 2, Table S5). Plasma TG content was 62-75% lower in starved fish than fed fish and was affected by a significant interaction between temperature and feeding status (F_(1,36)_=9.57, p=0.004; Table 2, Table S5).

### Experiment 2: Examining energy savings over winter in wild anadromous Arctic char

*Ṁ*O_2rest_ of wild anadromous Arctic char was 27% higher at the end of summer compared with the end of winter (F_(1,13)_=10.23, p<0.01; Figure 4a). *Ṁ*O_2max_ and AAS did not differ between the end of winter and summer (*Ṁ*O_2max_: F_(1,13)_=0.08, p=0.785; AAS: F_(1,13)_=0.23, p=0.638; Figure 4a).

**Figure 4.**
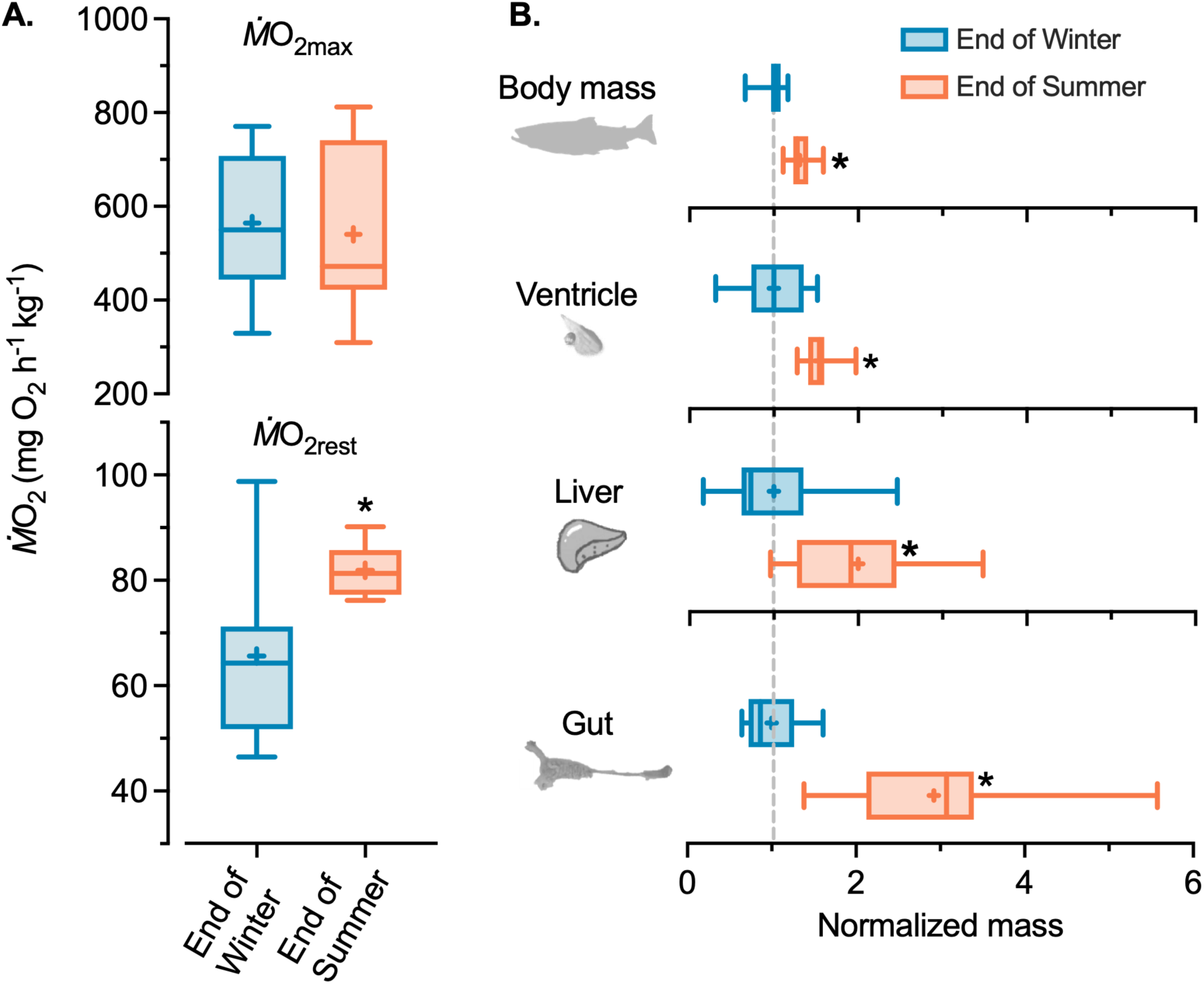
Seasonal resting and maximum rates of oxygen consumption (*Ṁ*O_2rest_ and *Ṁ*O_2max_; A.) in wild anadromous Arctic char (Experiment 2) with corresponding changes in normalized body and organ masses. Fish at ‘End of Winter’ had not started feeding or had only fed for up to ∼one week following winter fasting and fish at ‘End of Summer’ had presumably been feeding in the marine environment for the previous four to six weeks. Box-plots show median and interquartile range with 5-95% percentile whiskers for oxygen consumption (A.; n=8) adjusted for allometry to the average fish mass and body and organ masses (B.; n=10-31; Table S3) adjusted to average fish length and normalized as a proportion of the winter average (dashed line). An asterisk (*) represents a significant difference between end of winter and summer as assessed by GLM.

Fulton’s condition factor (K) was 28% greater at the end of summer compared with the end of winter (*X*^2^_(1)_=59.52, p<0.001; Table 3). After accounting for variation associated with body length, body mass and the masses of the ventricle, liver, stomach, pylorus, intestine and total gut were significantly greater at the end of summer compared with the end of winter (body mass: +27%, F_(1,41)_=53.63, p<0.0001; ventricle: +54%, F_(1,38)_=31.25, p<0.0001; liver: +100%, *X*^2^_(1)_=18.55, p<0.0001; stomach: +49%, *X*^2^_(1)_=23.09, p<0.0001; pylorus: +272%, *X*^2^_(1)_=88.12, p<0.0001; intestine: +374%, F_(1,40)_=58.19, p<0.0001; total gut: +192%, *X*^2^_(1)_=91.93, p<0.0001; Figure 4b). Given that the difference was less pronounced in the ventricle the gut, the gut:heart ratio was substantially higher at the end of summer than the end of winter (28.0 ± 2.7 vs. 44.2 ± 1.9, *X*^2^_(1)_=13.42, p<0.001). Relative intestine length did not differ between time points (F_(1,38)_=0.03, p=0.856).

**Table 3.** Size and condition of wild anadromous Arctic char caught at the end of winter or at the end of summer. End of winter represents fish that had recently migrated to sea and had either not resumed feeding or had only fed for a short amount of time (∼1 week), while end of summer represents fish that had fed for ∼ 4-6 weeks and were preparing to return to freshwater.

| **Respirometry** | **End of winter**<br>**(n=8)** | **End of summer**<br>**(n=8)** |
| --- | --- | --- |
| **Mass (kg)** | 4.24 ± 0.51 | 4.18 ± 1.03 |
| **Length (mm)** | 757.63 ± 42.76 | 701.25 ± 64.31 |
| **Fulton's Condition Factor** | 0.98 ± 0.08 | 1.20 ± 0.06* |
| **All Fish** | **(n=15)** | **(n=31)** |
| **Mass (kg)** | 3.74 ± 0.91 | 3.61 ± 1.45 |
| **Length (mm)** | 720.87 ± 65.75 | 651.23 ± 103.05 |
| **Fulton's Condition Factor** | 0.98 ± 0.08 | 1.24 ± 0.13* |

Data are means ± S.D. Significant differences from end of winter for Fulton's condition factor are indicated by an asterisk (\*).

Energy density was not different between the end of winter and summer (*X*^2^_(1)_=3.62, p=0.05; Table 4). The TG content of the gut (+179%; F_(1,21)_=132.6, p<0.0001) and white muscle (+140%; *X*^2^_(1)_=14.36, p<0.001) and total relative gut TG content (+580%; *X*^2^_(1)_=104.14, p<0.0001) were markedly higher at the end of summer (Table 4). Gut protein content was 37% lower at the end of summer (*X*^2^_(1)_=9.05, p<0.01), but total relative gut protein content was higher (F_(1,41)_=13.019, p<0.001; Table 4). White muscle protein content was 9% higher at the end of summer (*X*^2^_(1)_=4.25, p=0.045; Table 4).

**Table 4.** Energetic parameters of wild anadromous Arctic char captured at the end of winter or at the end of summer. End of winter represents fish that had recently migrated to sea and had either not started to feed or had only fed for a short amount of time (∼1 week), while end of summer represents fish that had fed for ∼ 4-6 weeks and were preparing to return to freshwater. Refer to Table S3 for exact sample sizes.

|  | **End of Winter**<br>**(n=13-20)** | **End of Summer**<br>**(n=10-31)** |
| --- | --- | --- |
| **Energy Density (kJ g dry mass<sup>-1</sup>)<sup>^</sup>** | 22.17 ± 0.31 | 23.31 ± 0.52 |
| **Gut TG Content (mg g<sup>-1</sup>)<sup>^</sup>** | 121.58 ± 13.72 | 339.39 ± 12.05* |
| **Total Relative Gut TG Content**<br>**(mg g body mass<sup>-1</sup>)** | 3.92 ± 0.57 | 25.81 ± 2.07* |
| **White Muscle TG Content (mg g<sup>-1</sup>)<sup>^</sup>** | 35.20 ± 5.39 | 84.31 ± 11.64* |
| **Gut Protein Content (mg g<sup>-1</sup>)<sup>^</sup>** | 257.33 ± 39.90 | 169.40 ± 14.34* |
| **Total Relative Gut Protein Content**<br>**(mg g body mass<sup>-1</sup>)** | 7.85 ± 1.19 | 13.08 ± 0.80* |
| **White Muscle Protein Content (mg g<sup>-1</sup>)<sup>^</sup>** | 697.79 ± 27.61 | 762.88 ± 18.10* |

Data are means ± S.E.M. An asterisk (\*) represents a significant difference from end of winter.
<sup>^</sup>Contents are per unit dry mass.
TG: Triglyceride

## Discussion

Species that face periods of ‘feast and famine’, such as the Arctic char, must balance the energetic challenge of food deprivation against the cost of maintaining physiological systems to support the daily rigours of life and maximize uptake when food becomes available (Armstrong & Schindler, 2011) (Armstrong and Bond 2013). By combining Arctic field-, and laboratory experiments we demonstrate that Arctic char suppress resting metabolic costs by selectively downregulating energetically expensive processes (i.e., protein synthesis and digestive tissue maintenance) that are extraneous during food deprivation. Wild anadromous Arctic char appear to do this each winter as they markedly increase resting metabolic demand, organ masses, fat stores, and body condition while feeding over summer. Thus, the mechanisms allowing energy conservation over winter are reversible on a timescale that does not compromise their ability to capitalize on summer food pulses. These responses are likely central to achieving a positive annual energy budget to support growth and reproduction despite the very brief summer foraging and energy acquisition window. That voluntary winter fasting in wild fish elicited similar physiological plasticity to that driven by forced starvation in the laboratory strengthens support for their ecological significance, demonstrating the value of better integrating field and laboratory approaches to understand animal energetics.

Another key finding was that maximal *Ṁ*O_2_ was not compromised with food deprivation despite reductions in resting *Ṁ*O_2,_ demonstrating that while digestive capacity was compromised, energy conservation responses need not trade-off against whole-animal metabolic performance. Previous studies in fishes and other species have provided evidence for this uncoupling of maximum and resting *Ṁ*O_2_ (Middleton et al. 2024) but its natural occurrence in the wild is not well documented in fish. Combined, these findings demonstrate the potential for fishes to express marked metabolic plasticity to cope with temporal variation in food availability while simultaneously maintaining key aspects of physiological performance to support necessary activities such as spring migrations and early foraging during the essential summertime re-accumulation of energy stores.

### Growth, and aerobic metabolism

Lowered resting metabolic demand is beneficial to slow the depletion of energy stores during periods of food limitation (Wang et al., 2006). Such reductions are likely essential in fishes like wild anadromous Arctic char that do not, or cannot, replenish energy stores over winter. Here, Arctic char substantially lowered resting metabolic demands with low temperature and food deprivation in the laboratory and in the wild in a similar manner to other salmonids (Auer, Salin, Rudolf, et al., 2016; Hvas et al., 2020; O’Connor et al., 2000; Van Leeuwen et al., 2012) (Middleton et al. 2024). In the laboratory, *Ṁ*O_2standard_ was more than halved with the initial cooling from 8°C to 2°C, reflecting the passive acute effect of cold on metabolic rate (Clarke & Johnston, 1999; Reeve et al., 2022). While fed fish at 2°C showed an impressive compensatory increase *Ṁ*O_2standard_ after 120 days of cold acclimation, *Ṁ*O_2standard_ remained below that of fed fish at 8°C, demonstrating the passive energy savings driven by cold alone. In contrast, the *Ṁ*O_2standard_ of starved fish at 2°C did not change over time despite drastic reductions in *K_s_* and tissue atrophy, which would inherently depress resting oxygen consumption as in starved fish at 8°C. Thus, starved Arctic char at 2°C were likely simultaneously increasing aspects of metabolism to compensate for cold limitations while down-regulating others to conserve energy in response to food deprivation. The result was that in all instances, wild and laboratory Arctic char had lower resting metabolic demands following prolonged food deprivation. Congeneric brook char (*Salvelinus fontinalis*) respond similarly in a laboratory setting to the combined effects of cold and food deprivation (Middleton et al., 2024). Together these results support our hypothesis that cold and food deprivation independently drive reductions in resting energy expenditure that are likely key to energy management when such conditions co-occur over winter.

Importantly, our field-based results likely represent conservative estimates of the scope for reductions in resting metabolic demand over winter. Ongoing acoustic telemetry studies, suggest that the wild Arctic char were likely in the ocean for approximately one week prior to capture (L.N. Harris, unpublished data), during which time they may have resumed feeding. This potential refeeding and the fact that *Ṁ*O_2rest_ may have been elevated in association with winter cold acclimation (as in Experiment 1) likely means that the difference in *Ṁ*O_2rest_ between seasons does not fully reflect the extent to which wild fish can reduce resting metabolic demand during winter fasting.

The maintenance of high aerobic metabolic performance has physiological and ecological implications for performance following periods of food deprivation (e.g., spring migration, and avoiding predation;) and the ability to capitalize on summer resource pulses (Auer et al., 2015). Maintaining *Ṁ*O_2max_ and aerobic scope with food deprivation, has been reported in other salmonids such as rainbow trout (*Oncorhynchus mykiss*), Atlantic salmon (*Salmo salar*) and chinook salmon (*O. tshawytscha*) (Alsop & Wood, 1997; Hvas, 2022; Thorarensen & Farrell, 2006). However, the duration of food deprivation in nearly all relevant studies has not exceeded one month and they do not explicitly study wild fish or natural food deprivation. Here, we show that following 120 days of starvation in the laboratory and that following prolonged winter food deprivation in the wild, Arctic char conserved *Ṁ*O_2max_ and aerobic scope while simultaneously reducing resting metabolic demands. Congruently, aspects of systems that support peak aerobic performance, such as plasma glucose content, blood-oxygen carrying capacity (i.e., hematocrit) and relative ventricular mass were also largely maintained with food-deprivation (Gallaugher et al., 1995; Ressel et al., 2021; Secor & Carey, 2016). In particular, the heart is a central mediator of *Ṁ*O_2max_ and can increase in mass with digestive capacity during refeeding in Dolly Varden (*Salvelinus malma*) (+35%; Armstrong & Bond, 2013) and Burmese pythons (*Python bivittatus*) (+40%; Secor, 2008), which also possess ‘feast and famine’ type life-histories. Likewise, relative to body length, wild Arctic char increased ventricular mass in proportion to body mass (i.e. similar heart-to-body mass ratio between seasons) likely supporting the similar mass specific *Ṁ*O_2max_ measured in both seasons.

### Tissue-specific responses to reductions in energy acquisition

Selective tissue atrophy is seen in many fishes (Armstrong & Bond, 2013; Jobling et al., 1998; Jørgensen et al., 1997; Martin et al., 1993), amphibians (Naya et al., 2009), reptiles (Secor, 2008; Secor et al., 1994), birds (Funes et al., 2014) and mammals (Fedorov et al., 2009) in response to food deprivation. For example, other salmonids such as Dolly Varden and Atlantic salmon also show significant atrophy of the gut (-49-154%) with food deprivation related to pulse resource use and anadromy (Armstrong & Bond, 2013; Martin et al., 1993). The effects on energy expenditure, however, have not been measured. Here, food deprivation in Arctic char resulted in loss of organ mass for most organs examined; but was largely focused in the gut, where continuous maintenance is likely inefficient with predictable and prolonged intervals between feeding (Armstrong & Schindler, 2011; Secor, 2008). The extent of atrophy observed in our field study, while impressive, is likely an underestimate of true winter atrophy in anadromous Arctic char as discussed for *Ṁ*O_2rest_. Nonetheless, in field and laboratory assessments, substantial organ-specific reductions in mass were associated with reductions in resting *Ṁ*O_2_ and thus energy expenditure.

While resting metabolic demand was markedly reduced in the absence of food, it was not sufficient to avoid a substantial loss of energy stores. Food deprivation in the laboratory resulted in large reductions in body fat, energy density, visceral and muscle TG content, and increased moisture content as expected (Brett et al., 1969; Cassidy et al., 2016; Jobling et al., 1998; Navarro & Gutierrez, 1995; Secor & Carey, 2016). Yet protein content was conserved to a greater extent suggesting a robust starvation tolerance as recently demonstrated in brook char (Middleton et al. 2024). Wild anadromous Arctic char showed significant loss of TG content in the gut and white muscle following ∼10 months of food deprivation over winter but no change in energy density of the white muscle, highlighting the use of visceral fat as a primary transient fat source (Navarro & Gutierrez, 1995). In the field, the loss of protein was more substantial than in the laboratory which probably reflects the much longer duration of food deprivation (Navarro & Gutierrez, 1995; Shuter et al., 2012). Thus, wild anadromous Arctic char may be in the critical phase of starvation (i.e., protein catabolism) at the end of winter with refeeding necessary to avoid mortality (Secor & Carey, 2016).

### Adjustments of in vivo protein synthesis rates as an underlying mechanism for energy savings

The synthesis of new proteins is a significant contributor to energy expenditure in fishes, accounting for ∼20-50% of resting energy demands (Carter & Houlihan, 2001). This association is highlighted by the strong positive correlations we found between *K_s_* and *Ṁ*O_2standard_. Reducing *K_s_* is a commonly reported response to food deprivation among salmonids including rainbow trout, brook char and Arctic char, however, past studies have considered shorter durations (Middleton et al. 2024)(Cassidy et al., 2016; Loughna & Goldspink, 1984; McMillan & Houlihan, 1992; Smith, 1981) (Middleton et al. 2024). Here, we found moderate reductions in *K_s_* in the ventricle, and liver, but substantial reductions in gut and white muscle following ∼140 days of starvation. Additionally, reductions in liver and gut *K_s_* co-occurred with a loss of tissue mass amplifying whole-organ energy savings. Decreases in routine levels of gene expression, ATPase enzyme activities and mitochondrial activity during food deprivation (Cassidy et al., 2016; Méndez & Wieser, 1993; Ressel et al., 2021; Salin et al., 2018; Secor & Carey, 2016; Thoral et al., 2023) are among other strategies that may provide further energy savings and warrant further study.

### Conclusions and ecological implications

Animals faced with sustained environmental challenges, including the frigid temperatures and low food availability characteristic of high-latitude winters, commonly express varying degrees of dormancy to circumvent associated performance or energy decrements (Ultsch 1989, Shuter et al. 2012, Reeve et al. 2022). Indeed, overwintering strategies in fishes can be characterized on a spectrum from winter dormant to lethargic to active (i.e., non-dormant) based on the degree to which they reduce feeding, activity, and metabolism (Shuter et al. 2012, Reeve et al. 2022). While Arctic char have the physiological ability to remain active and feed at near zero temperatures (Gilbert, 2020; Dubos et al. Under Review), they nonetheless exhibit an integrated overwintering strategy with multiple hallmarks of winter dormancy: 1) Arctic char commonly exhibit a programmed (i.e., not strictly temperature dependent) winter fasting in the wild, and laboratory setting despite continued food availability (Tveiten et al., 1996) (Jorgensen and Johnsen 2014) 2) They markedly reduce or cease activity for prolonged periods, and do so to a much greater extent than sympatric species that remain feeding overwinter (Dubos et al, 2025) and 3) As detailed here, Arctic char further reduce energy expenditure passively as a result of cooling, and actively through reductions in organ volume and protein synthesis rates that are a product of food deprivation (i.e. not cold-induced metabolic depression). These same hallmarks are characteristic of winter dormancy in terrestrial animals. Thus, the present research on Arctic char, and previous work on overwintering in fish (Reeve et al, 2022; Shuter 2012, Ultsch 1989), demonstrate that a unifying framework for dormancy recently developed primarily for terrestrial animals (Wilsterman et al. 2021) can and should be expanded to the aquatic realm.

Such dormancy demonstrates that animals that experience ‘feast and famine’ can optimize their energy budgets by adjusting energy expenditure in concert with food availability through physiological and behavioural plasticity (Auer, Salin, Anderson, et al., 2016; Hvas et al., 2020; Naya et al., 2009; O’Connor et al., 2000; Tøien et al., 2011; Van Leeuwen et al., 2012).

From an energetic perspective, such plasticity, however, is only beneficial if the energy saved is greater than the associated direct (i.e., remodelling) and indirect (e.g. forgone feeding opportunity) costs. For instance, Armstrong and Schindler (2011), found that despite the costly nature of digestive machinery, predatory fishes commonly maintain greater digestive capacity than they consistently use, to ensure that they can capitalize on patchy, intermittent feeding opportunities. In contrast, when food is restricted on a prolonged or predictable basis, as in the present study, the balance shifts in favor of strategies to suppress metabolism including those that compromise digestive capacity. Thus, examinations of fish bioenergetics need to consider that metabolism varies as fish balance the trade-offs between the costs of plasticity and maintaining physiological systems (Armstrong and Schindler 2011). Lastly, further consideration should be given to how climate change will impact these energy management strategies; warmer temperatures and changes in the timing of seasonal events can have pronounced direct and indirect effects on fish metabolism and patterns of food availability thereby shaping the viability of such strategies.

## Acknowledgements

The authors thank the Ekaluktutiak Hunters and Trappers Organization and members of the community of Ikaluktutiak for their support. We thank Beverly Maksagak for helping coordinate field research, Kevin Kanayok for his contribution as a field technician, and Dr. Bonnie Hamilton who contributed to field sampling in 2019. Laboratory studies were approved by the Animal Care Committee of the University of New Brunswick (UNB), Saint John, (UNB 2020-4S-03) and field sampling and animal use was approved by Fisheries and Oceans Canada Freshwater Institute (AUP-2019-37, AUP-2021-48, AUP-2022-82, AUP-2023-61, LFSP S-19/20-1017-NU, LFSP S-21/22-1021-NU, LFSP S-22/23-1043-NU).

## Data Availability

All supporting data will be archived on FigShare upon publication

