## Supplemental Material for "Metabolic plasticity facilitates a high-latitude life of feast and famine in Arctic char"

**Supplemental Information**

**Experiment 1: The effects of prolonged starvation at cold temperatures on the energetics of juvenile Arctic char in a laboratory setting**

*Validation of* $\dot{M}O_{2standard}$ *estimates*

Estimates of $\dot{M}O_{2standard}$ can be influenced by the level of activity of fish within the respirometer. To estimate $\dot{M}O_{2standard}$ at zero activity we generated estimated marginal means at zero activity using GLMM (family=Gamma, link=log) with $\dot{M}O_2$ ($mgO_2\ h^{-1}$) as a function of activity, temperature, and feeding status with body mass included as a covariate and fish ID included as a random factor. We took the antilog of the estimated marginal means of $\dot{M}O_{2standard}$ and its standard error to compare to the values obtained through the traditional 20$^{th}$ percentile method (Table S1). We calculated $\dot{M}O_{2standard}$ using the 20$^{th}$ percentile method at 120 days using all fish and using only fed fish that grew, but we included all fish in the calculation of the estimates at zero activity using the GLMM (Table S1).

*Organ protein content*

Protein content of the liver and white muscle and total relative gut protein content did not differ between treatment groups (Table 3). Total relative liver protein content was affected by an interaction between temperature and feeding status ($F_{(1,34)}=6.93$, $p<0.05$) that arose because temperature only had an impact if the fish were fed (Table 3). Fed fish had significantly higher (2°C: +94%; 8°C: +63%) total relative liver protein content than starved fish. Starved fish at 2°C had significantly higher gut protein content than both groups of fed fish ($p<0.01$; Table 3).

**Tables and Figures**

**Table S1.** Validation of $\dot{M}O_{2standard}$ estimates from juvenile Arctic char that were fed daily (0.5% body mass ration) or starved at 8°C or 2°C for 120 days. The $\dot{M}O_{2standard}$ was calculated using the 20th percentile method (all fish and fed fish that grew only) and generated estimated marginal means at zero activity (GLMM; Family=Gamma, link=log).

| | $\dot{M}O_{2standard}$ obtained through 20th percentile method (all fish) | $\dot{M}O_{2standard}$ obtained through 20th percentile method (fed growers only) | $\dot{M}O_{2standard}$ estimated at zero activity with a GLMM |
|---|---|---|---|
| **Fed at 2°C** | 27.49 ± 7.51 | 30.56 ± 6.66 | 27.80 ± 6.83 |
| **Fed at 8°C** | 41.26 ± 7.58 | 46.90 ± 6.63 | 48.67 ± 6.88 |
| **Starved at 2°C** | 15.91 ± 7.53 | 15.78 ± 6.62 | 17.55 ± 6.81 |
| **Starved at 8°C** | 26.65 ± 7.59 | 26.56 ± 6.71 | 27.53 ± 6.88 |

$\dot{M}O_{2standard}$, Standard rate of oxygen consumption
GLMM, Generalized linear mixed effects model

**Table S2.** Thermal sensitivity quotients ($Q_{10}$) of oxygen consumption rates, aerobic scope and protein synthesis rates of Arctic char. $Q_{10}$'s were calculated using the mean values at each temperature.

| Trait | Temperature Interval | Time Period (days) | Fed | Starved |
|---|---|---|---|---|
| $\dot{M}O_{2standard}$ ($mgO_2$ $h^{-1}$ $kg^{-1}$) | 8°C – 2°C | 0 | 3.5 | 3.3 |
| | | 30 | 3.2* | 2.2 |
| | | 60 | 3.0* | 1.9 |
| | | 120 | 2.0 | 2.4 |
| $\dot{M}O_{2max}$ ($mgO_2$ $h^{-1}$ $kg^{-1}$) | 8°C – 2°C | 120 | 1.8 | 1.2 |
| Absolute Aerobic Scope ($mgO_2$ $h^{-1}$ $kg^{-1}$) | 8°C – 2°C | 120 | 1.8 | 1.1 |
| Fractional Protein Synthesis Rates ($K_s$; % $day^{-1}$) | 8°C – 2°C | Liver: 142 | 2.3 | 2.2 |
| | | Gut: 142 | 2.0 | 1.8 |
| | | Muscle: 142 | 0.6 | 1.0 |
| | | Ventricle: 142 | 2.7 | 2.6 |

$\dot{M}O_{2standard}$, Standard rate of oxygen consumption
$\dot{M}O_{2max}$, Maximum rate of oxygen consumption
*Several fish at 2°C were volitionally fasting potentially masking acclimation responses

**Table S3.** Summary of the sample sizes for field measurements of wild anadromous Arctic char at the end of winter and the end of summer. Sample sizes may differ from Table 1 because we did not have mass and length for every individual used in energetic analyses.

| | End of Winter | End of Summer |
|---|---|---|
| **Oxygen Consumption Rates ($\dot{M}O_2$)** | n=8 | n=8 |
| **Ventricle Mass** | n=13 | n=28 |
| **Liver Mass** | n=13 | n=31 |
| **Stomach, Pylorus and Intestine and Gut Mass** | n=13 | n=30 |
| **Intestine Length** | n=11 | n=30 |
| **Gut:Heart** | n=11 | n=27 |
| **Energy Density** | n=20 | n=20 |
| **Gut Triglyceride** | n=13 | n=10 |
| **Total Relative Gut Triglyceride** | n=13 | n=10 |
| **White Muscle Triglyceride** | n=20 | n=10 |
| **Gut Protein** | n=13 | n=29 |
| **Total Relative Gut Protein** | n=13 | n=29 |
| **White Muscle Protein** | n=20 | n=31 |

**Table S4. (See next page)** Summary of statistical outputs obtained from generalized linear mixed effects models (GLMM; family=Gamma, link=log) or linear mixed effects models (LMM; median activity) examining the effects of temperature (2°C or 8°C), prolonged starvation, time and their interactions on the morphometrics and standard oxygen consumption rates ($\dot{M}O_{2standard}$) of juvenile Arctic char (Experiment 1). These analyses correspond with data shown in Figure 2 and Table 2. Significance ($p<0.05$) was assessed using type II Wald chi-square tests and is indicated in bold. Length, mass and Fulton's condition factor were included only for fish that had complete datasets (i.e., mass and length were measured at each time point).

| | **Temperature** (df=1) | | **Feeding Status** (df=1) | | **Time** (df=3) | | **Body Mass** (df=1) | | **Temperature: Treatment** (df=1) | | **Temperature: Time** (df=3) | | **Treatment: Time** (df=3) | | **3-way Interaction** (df = 3) | |
|---|---|---|---|---|---|---|---|---|---|---|---|---|---|---|---|---|
| | $X^2$ | p | $X^2$ | p | $X^2$ | p | $X^2$ | p | $X^2$ | p | $X^2$ | p | $X^2$ | p | $X^2$ | p |
| **Length** | 0.47 | 0.49 | 1.69 | 0.19 | 115.79 | **<0.01** | | | 0.17 | 0.68 | 40.30 | **<0.01** | 157.78 | **<0.01** | 64.05 | **<0.01** |
| **Body Mass** | 0.07 | 0.79 | 5.0 | 0.03 | 100.08 | **<0.01** | | | 0.59 | 0.44 | 29.70 | **<0.01** | 317.19 | **<0.01** | 78.58 | **<0.01** |
| **Body Condition** | 0.85 | 0.36 | 8.25 | **<0.01** | 7.49 | 0.06 | | | 1.54 | 0.21 | 7.92 | **0.048** | 156.54 | **<0.01** | 15.20 | **<0.01** |
| **$\dot{M}O_{2standard}$** | 65.75 | **<0.01** | 6.83 | **<0.01** | 7.03 | 0.07 | 62.96 | **<0.01** | 0.45 | 0.50 | 16.96 | **<0.01** | 43.17 | **<0.01** | 10.65 | **0.01** |

**Table S5.** Summary of statistical outputs obtained from linear models (LM) examining the effects of daily feeding (0.5% body mass ration) or prolonged starvation for 120 days at 2°C or 8°C and their interactions on various performance variables of juvenile Arctic char. These analyses correspond with the data shown in Figures 2, 3 and 4 and Table 3 in the main text. Significant effects ($p<0.05$) were calculated using type II Wald chi-square tests and are indicated in bold.

| | Temperature | | | Feeding Status | | | Body Length* or Body Mass^ | | | Interaction | | |
|---|---|---|---|---|---|---|---|---|---|---|---|---|
| | F | df | p | F | df | p | F | df | p | F | df | p |
| **$\dot{M}O_{2standard}$^** | 57.63 | 1,35 | **<0.01** | 59.58 | 1,35 | **<0.01** | 40.42 | 1,35 | **<0.01** | 0.50 | 1,35 | 0.49 |
| **Body Fat** | 0.89 | 1,36 | 0.35 | 49.58 | 1,36 | **<0.01** | | | | 1.59 | 1,36 | 0.21 |
| **Moisture Content** | 0.07 | 1,36 | 0.79 | 16.23 | 1,36 | **<0.01** | | | | 0.69 | 1,36 | 0.42 |
| **Energy Density** | 0.07 | 1,35 | 0.79 | 29.38 | 1,35 | **<0.01** | | | | 2.19 | 1,35 | 0.15 |
| **Hematocrit** | 2.0 | 1,36 | 0.17 | 1.71 | 1,36 | 0.19 | | | | 6.48 | 1,36 | **0.02** |
| **Plasma TG** | 11.64 | 1,36 | **<0.01** | 71.29 | 1,36 | **<0.01** | | | | 9.57 | 1,36 | **<0.01** |
| **Ventricle Mass^** | 2.89 | 1,35 | 0.097 | 0.68 | 1,35 | 0.42 | 317.8 | 1,35 | **<0.01** | 1.69 | 1,35 | 0.201 |
| **Liver Mass^** | 2.85 | 1,35 | 0.10 | 39.19 | 1,35 | **<0.01** | 179.56 | 1,35 | **<0.01** | 5.93 | 1,35 | **0.02** |
| **Pylorus Mass^** | 0.30 | 1,35 | 0.59 | 35.25 | 1,35 | **<0.01** | 116.64 | 1,35 | **<0.01** | 2.35 | 1,35 | 0.13 |
| **Intestine Length*** | 4.99 | 1,34 | **0.03** | 61.32 | 1,34 | **<0.01** | 25.72 | 1,34 | **<0.01** | 2.47 | 1,34 | 0.13 |
| **Gut Mass^** | 0 | 1,35 | 0.99 | 28.96 | 1,35 | **<0.01** | 115.03 | 1,35 | **<0.01** | 0.55 | 1,35 | 0.46 |
| **Liver $K_s$** | 44.25 | 1,36 | **<0.01** | 23.90 | 1,36 | **<0.01** | | | | 2.12 | 1,36 | 0.15 |
| **Carcass Protein Content^** | 0.04 | 1,35 | 0.83 | 3.56 | 1,35 | 0.067 | 1.69 | 1,35 | 0.20 | 0.07 | 1,35 | 0.80 |
| **Liver Protein Content** | 0.14 | 1,35 | 0.71 | 0.27 | 1,35 | 0.61 | | | | 3.78 | 1,35 | 0.06 |
| **Total Relative Liver Protein Content^** | 8.01 | 1,34 | **<0.01** | 47.73 | 1,34 | **<0.01** | 2.45 | 1,34 | 0.13 | 6.93 | 1,34 | **0.01** |
| **Gut Protein Content** | 2.09 | 1,32 | 0.16 | 18.46 | 1,32 | **<0.01** | | | | 0.57 | 1,32 | 0.46 |
| **Total Relative Gut Protein Content^** | 4.83 | 1,31 | **0.04** | 1.99 | 1,31 | 0.17 | 2.97 | 1,31 | 0.09 | 0.72 | 1,31 | 0.40 |
| **White Muscle Protein Content** | 0.34 | 1,31 | 0.56 | 8.91 | 1,31 | **<0.01** | | | | 0.02 | 1,31 | 0.89 |

$\dot{M}O_{2standard}$, Standard oxygen consumption
TG, Triglyceride
$K_s$, Protein synthesis rate

**Table S6.** Summary of statistical outputs obtained from generalized linear models (GLM; family=Gamma, link=log) examining the effects of daily feeding (0.5% body mass ration) or prolonged starvation for 120 days at 2°C or 8°C and their interactions on various performance variables of juvenile Arctic char. These analyses correspond with the data shown in Figures 2, 3 and 4 and Table 3 in the main text. Significant effects ($p<0.05$) were calculated using type II Wald chi-square tests and are indicated in bold.

| | **Temperature** (df=1) | | **Feeding Status** (df=1) | | **Body Mass** (df=1) | | **Temperature: Treatment** (df=1) | |
|---|---|---|---|---|---|---|---|---|
| | $X^2$ | p | $X^2$ | p | $X^2$ | p | $X^2$ | p |
| **$\dot{M}O_{2max}$** | 13.95 | **<0.001** | 0.30 | 0.58 | 46.52 | **<0.001** | 3.32 | 0.07 |
| **AAS** | 8.43 | **0.004** | 0.29 | 0.59 | 36.21 | **<0.001** | 3.13 | 0.08 |
| **Stomach Mass** | 0.69 | 0.41 | 44.43 | **<0.001** | 47.45 | **<0.001** | 9.60 | **0.002** |
| **Intestine Mass** | 3.06 | 0.08 | 127.24 | **<0.001** | 51.66 | **<0.001** | 0.29 | 0.59 |
| **Gut:Heart Ratio** | 0.07 | 0.80 | 91.22 | **<0.001** | 0.03 | 0.87 | 1.03 | 0.31 |
| **Glucose** | 4.65 | **0.03** | 2.39 | 0.12 | | | 1.16 | 0.28 |
| **Gut TG Content** | 1.43 | 0.23 | 19.74 | **<0.001** | | | 0.96 | 0.33 |
| **Total Relative Gut TG** | 1.25 | 0.26 | 12.16 | **<0.001** | 3.94 | **0.047** | 0.39 | 0.53 |
| **White Muscle TG Content** | 4.19 | **0.04** | 17.97 | **<0.001** | | | 0.36 | 0.55 |
| **Ventricle $K_s$** | 66.52 | **<0.001** | 43.68 | **<0.001** | | | 0.007 | 0.93 |
| **Gut $K_s$** | 18.68 | **<0.001** | 153.78 | **<0.001** | | | 0.04 | 0.83 |
| **White Muscle $K_s$** | 1.92 | 0.17 | 96.85 | **<0.001** | | | 2.41 | 0.12 |

$\dot{M}O_{2max}$, Maximum oxygen consumption
AAS, Absolute aerobic scope
TG, Triglyceride
$K_s$, Protein synthesis rate

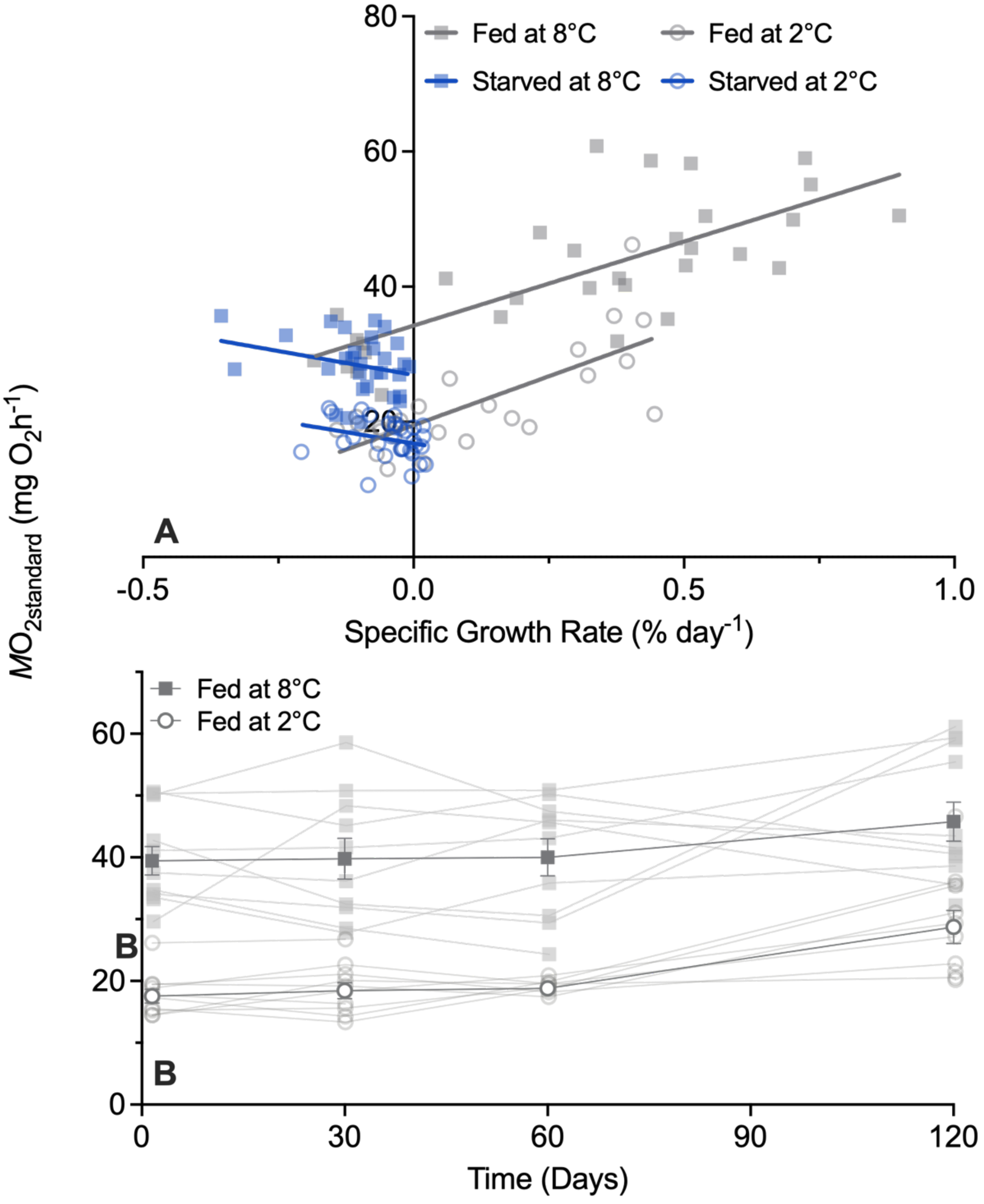


**Figure S1.** Linear regression between (A) $\dot{M}O_{2standard}$ and specific growth rate (% day$^{-1}$) and (B) the standard rate of oxygen consumption ($\dot{M}O_{2standard}$; mg$O_2$ h$^{-1}$ kg$^{-1}$) of fed individuals in juvenile Arctic char over 120 days of daily feeding (0.5% body mass ration; closed) or prolonged starvation (open) at 2°C (blue) or 8°C (red). For the regressions (A), data are included for measurements at 30, 60 and 120 days but exclude the 'day 0' measurement due to lack of growth information (i.e., ‘day 0’ was the first measurement). Linear regressions are shown for fish fed at 2°C ($R^2 = 0.55$, $p<0.0001$), starved fish at 2°C ($R^2 = 0.08$, $p>0.05$), fed fish at 8°C ($R^2 = 0.57$, $p<0.0001$) and starved fish at 8°C ($R^2 = 0.07$, $p>0.05$). For $\dot{M}O_{2standard}$ over time (B), data are shown for the mean ± S.E.M. for groups of fed fish at 2°C (blue circles) and 8°C (grey circles) along with all individual measurements. Feeding treatments started at ‘day 0’ and temperature treatment for 2°C groups started 24 hours prior to the ‘day 0’ measurement following acute cooling (~10 hours).

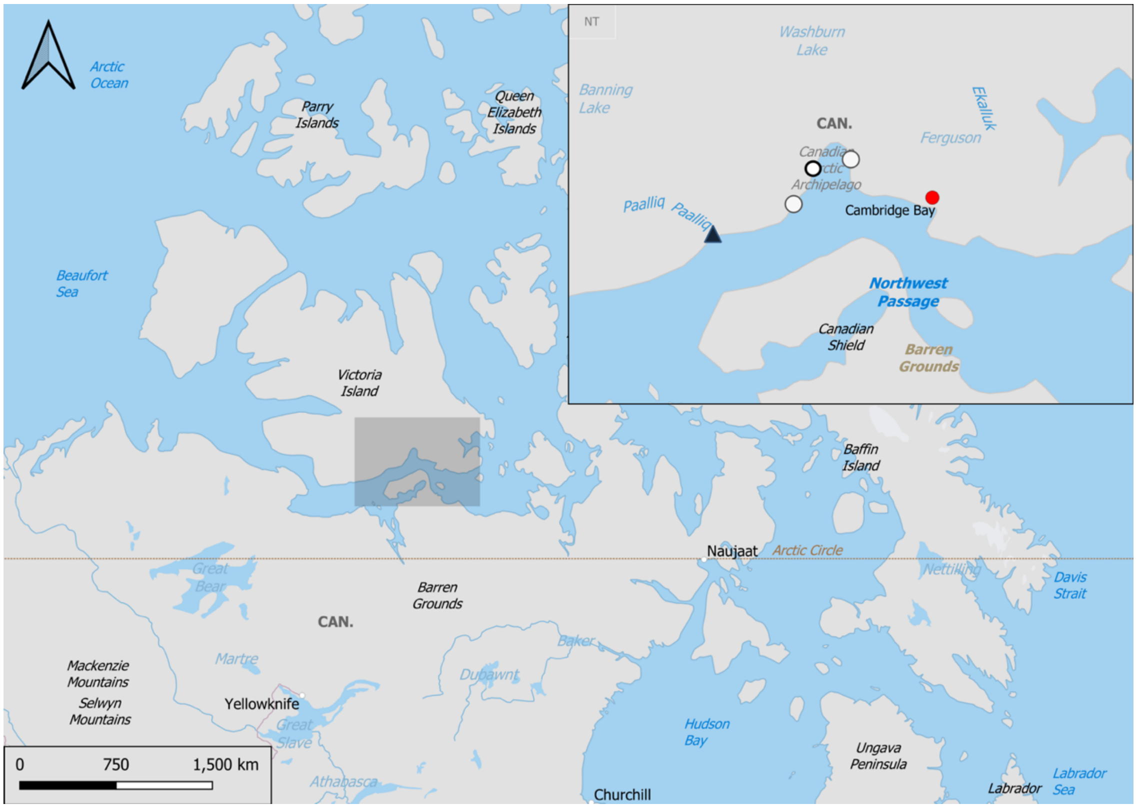


**Figure S2.** Study area in the Kitikmeot region of Nunavut, Canada. Anadromous Arctic char were captured at Palik, the mouth of the Lauchlan River as it drains into the Arctic Ocean. The inset shows the shaded region with the study site indicated by a black triangle; the nearby commercial fishery sites (Halokvik, Surrey and Ekalluktok) are represented by white circles and the community of Ikaluktutiak (Cambridge Bay) is indicated by a red circle.
